# MRI-based computational model generation for cerebral perfusion simulations in health and ischaemic stroke

**DOI:** 10.1101/2022.09.07.506940

**Authors:** T. I. Józsa, J. Petr, F. Barkhof, S. J. Payne, H. J. M. M. Mutsaerts

**Affiliations:** Department of Radiology and Nuclear Medicine, Amsterdam Neuroscience, Amsterdam University Medical Centers, Location VUmc, Amsterdam, the Netherlands; Institute of Biomedical Engineering, Department of Engineering Science, University of Oxford, Parks Road, Oxford OX1 3PJ, UK; Helmholtz-Zentrum Dresden-Rossendorf, Institute of Radiopharmaceutical Cancer Research, Dresden, Germany; UCL Queen Square Institute of Neurology, University College London, London, UK; Centre for Medical Image Computing (CMIC), Faculty of Engineering Science, University College London, London, UK; Institute of Applied Mechanics, National Taiwan University, Taipei, Taiwan

## Abstract

Cerebral perfusion models were found to be promising research tools to predict the impact of acute ischaemic stroke and related treatments on cerebral blood flow (CBF) linked to patients’ functional outcome. To provide insights relevant to clinical trials, perfusion simulations need to become suitable for group-level investigations, but computational studies to date have been limited to a few patient-specific cases. This study set out to overcome issues related to automated parameter inference, that restrict the sample size of perfusion simulations, by integrating neuroimaging data. Seventy-five brain models were generated using measurements from a cohort of 75 healthy elderly individuals to model resting-state CBF distributions. Computational perfusion model geometries were adjusted using healthy reference subjects’ T1-weighted MRI. Haemodynamic model parameters were determined from CBF measurements corresponding to arterial spin labelling perfusion MRI. Thereafter, perfusion simulations were conducted for 150 acute ischaemic stroke cases by simulating an occlusion and cessation of blood flow in the left and right middle cerebral arteries. The anatomical (geometrical) fitness of the brain models was evaluated by comparing the simulated grey and white matter (GM and WM) volumes to measurements in healthy reference subjects. Statistically significant, strong positive correlations were found in both cases (GM: Pearson’s *r* 0.74, *P*-value< 0.001; WM: Pearson’s *r* 0.84, *P*-value< 0.001). Haemodynamic parameter tuning was verified by comparing total volumetric blood flow rate to the brain in reference subjects and simulations resulting in Pearson’s *r* 0.89, and *P*-value< 0.001. In acute ischaemic stroke cases, the simulated infarct volume using a perfusion-based proxy was 197±25 ml. Computational results showed excellent agreement with anatomical and haemodynamic literature data corresponding to T1-weighted, T2-weighted, and phase-contrast MRI measurements both in healthy scenarios and in acute ischaemic stroke cases. Simulation results represented solely worst-case stroke scenarios with large infarcts because compensatory mechanisms, e.g. collaterals, were neglected. The established computational brain model generation framework provides a foundation for population-level cerebral perfusion simulations and for *in silico* clinical stroke trials which could assist in medical device and drug development.

## 1 Introduction

Acute ischaemic strokes (AIS) are caused by occlusions of major cerebral arteries and account for the majority of stroke cases [1, 2]. Thrombolysis [2] and thrombectomy [1] revolutionised stroke treatment but up to 66% of patients remain functionally dependent [3, 4]. The *IN Silico* clinical trials for the treatment of acute Ischaemic STroke (INSIST) consortium [5, 6] set out to improve existing stroke treatments based on computational models [7–11]. In AIS, perfusion shortage is the primary driver of infarct formation associated with irreversible brain tissue and function loss. Therefore, perfusion modelling is a key element of the INSIST simulation framework [5, 6].

In the context of stroke simulations, the purpose of computing cerebral perfusion is to estimate blood flow in healthy, occluded, and post-treatment states and to provide insights comparable to clinical measurements. To this end, models have been proposed describing microscale arterioles, capillaries, and venules embedded into the brain parenchyma as coupled porous media [12, 13]. Similar multi-compartment models have been utilised previously in a more generic poroelastic framework [14, 15]. These models require knowledge of anatomical (geometrical) and pathophysiological parameters, such as the volumetric meshes of reconstructed brain geometries [16, 17], and the permeability of the microcirculation [7, 18, 19]. The automated inference of such parameters remains an outstanding challenge. For this reason, studies on computational perfusion simulations have been limited to less than three brain models [12, 13, 15, 20]. Intervention trials incorporate typically hundreds of individuals to ensure the safety and efficacy of drugs and medical devices [1]. Therefore, the small sample size remains a major obstacle to the widespread application of perfusion simulations.

The present study sets out to overcome the limitation of the small number of computational brain models in perfusion simulations. The proposed method uses clinical measurements, including structural and perfusion-weighted magnetic resonance imaging (MRI), to generate automatically multiple models with representative perfusion distributions corresponding to healthy reference subjects’ resting states. Cohort-level perfusion modelling allows us to directly compare simulation results with clinical measurements in healthy individuals and in AIS patients.

## 2 Theory

This section describes the theoretical aspects of the proposed perfusion simulations. Firstly, a mathematical framework is presented and linked to anatomy and physiology through parametrisation. Thereafter, Section 3 covers how the necessary parameters can be obtained from clinical data including neuroimaging.

The employed perfusion model builds on previous results regarding two- [12] and three-compartment [13–15, 20] porous haemodynamics models. Parenchymal blood vessels are categorised as arterioles, capillaries, and venules. The microcirculation is described as two porous media – arteriole and venule vessel networks – using local “voxel-scale” space-averaged variables and macroscale parameters, such as the blood pressure and permeability tensors representing the conductance of descending arteriole and ascending venule vessel bundles. The arteriole and venule compartments are then connected by a coupling term symbolising the “voxel-scale” conductance of capillaries. The number of compartments in a multi-compartment porous formulation depends on the categorisation of microvessels determined by the context of use. Therefore, this approach does not require detailed knowledge of microscale blood vessels.

### 2.1 Mathematical formulation

The porous perfusion model utilised in the present study targets AIS modelling. Therefore, it focuses on the arterial side of the systemic cerebral circulation and accounts solely for a single arteriole compartment. This simplification can be derived directly from a two-compartment model [12, 20] by assuming constant venule pressure. Cerebral perfusion pressure (CPP) defined as the arteriole and venule pressure difference is the primary variable, with spatial distribution of CPP equal to:

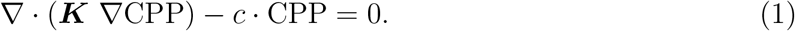

In this section, the International System of Units (SI) is used to ease dimensional analysis so that CPP [Pa = kgm^-1^ s^-2^]. In Equations (1), ∇ is the first-order spatial differential operator, so that ∇CPP [kgm^-2^ s^-2^] is the gradient of the perfusion pressure. The first term in Equation (1) represents fluid flow in arterioles as dictated by CPP and the permeability tensor ***K*** [m^3^ skg^-1^]. Darcy’s law characterises flow in porous media with the ratio of permeability [m^2^] and the dynamic viscosity [kgm^-1^ s^-1^] of the filling fluid. In the case of blood flow, viscosity depends on the pore size [21], which is not constant in the microcirculation. Therefore, these two quantities cannot be separated on a global level, and thus both of them are incorporated in the permeability tensor ***K*** [18, 19].

The second term in Equation (1) describes local drainage, which is linearly proportional to the arteriole-venule coupling coefficient, *c* [m s kg^-1^]. The *c* · CPP [s^-1^] term quantifies the volume of blood leaving the arteriole compartment locally in a unit time per given tissue volume. Divided by the brain tissue density *ρ* [kgm^-3^], it provides perfusion [m^3^ s^-1^ kg^-1^] – a term defined here interchangeably with cerebral blood flow (CBF) – as

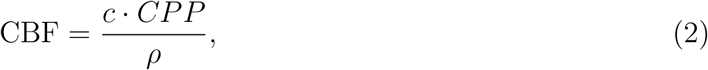

The model is quasi-steady-state and does not include gravity, and thus corresponds to shortterm time-averaged values in a lying individual’s resting state. We assume *ρ* = 1000 kgm^-3^ as estimates lie between 978 [22, 23] and 1040 kgm^-3^ [24].

### 2.2 Parametrisation

Here, the parametrisation of the perfusion model from the previous section is described summarizing our previous work [13, 20]. To capture rapid perfusion change between grey and white matter, different coupling coefficients are allocated for the corresponding tissue regions denoted by *c*^GM^ and *c*^WM^, respectively. Thereafter, a reference Cartesian coordinate system is defined to compute ***K***. The first axis of this reference coordinate system is aligned with the local centreline of descending arterioles. It is assumed herein that descending arterioles are not connected and therefore their permeability tensor in the reference coordinate system is

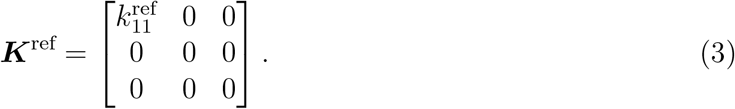

From here, the anisotropic and heterogeneous ***K*** tensor field is obtained by transforming ***K***^ref^ from the local (reference) to the global coordinate system as detailed in [13]. The reference coordinate system is computed from the brain geometry, assuming that: (i) penetrating arterioles descend perpendicularly to the cortical surface [25]; and (ii) arterioles follow the shortest path between the cortical and ventricular surfaces. Model parameters used in this study are summarised in Table 1.

**Table 1:**
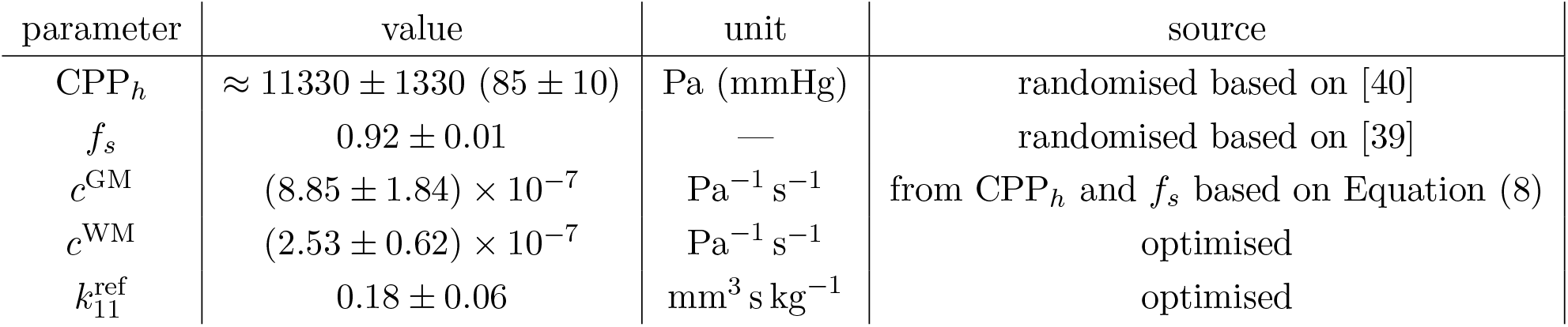
Haemodynamic parameters of the computational perfusion models simulated in this study (mean±SD). CPP - cerebral perfusion pressure, *f_s_* - CPP scaling factor, *c*^GM^ and *c*^WM^ - grey and white matter coupling coefficients, 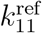 - arteriole permeability in a conveniently defined reference coordinate system.

### 2.3 Geometrical definition and boundary conditions

Equation (1) is solved over the domain of interest restricted to the brain tissue denoted by Ω, comprised of grey and white matter. The bounding surfaces (*∂*Ω) are

1. the ventricular surfaces Γ_*V*_; and
2. the pial surface Γ_*P*_,

so that *∂*Ω = Γ_*V*_ ∪ Γ_*P*_. The boundary region associated with the pial surface is divided further into eight subregions of the major feeding arteries using a vascular territory atlas [26]. The surface regions correspond to the left and right anterior, middle and posterior cerebral arteries (six regions), the cerebellar arteries (1 region), and the perforating basilar and vertebral arteries perfusing the brain stem (1 region). This subdivision is necessary to impose boundary conditions for stroke simulations associated with major cerebral artery occlusion.

The imposed boundary conditions are:

- Blood flow rate through the ventricle surfaces (Γ_*V*_) is zero in every case. Introducing ***n*** as the outward-pointing normal unit vector corresponding to the boundary surfaces, this Neumann type boundary condition reads as

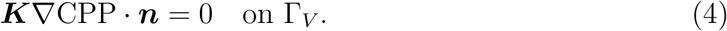 Filtration through the blood-cerebrospinal fluid barrier is neglected but it could be modelled by setting a non-zero value on the right-hand side of Equation (4) for Γ_*V*_ to represent fluid flux at the choroid plexus.
- The Γ_*P*_ boundary treatment distinguishes two different scenarios:
  – In healthy cases, constant cerebral perfusion pressure (CPP_h_) [20]:

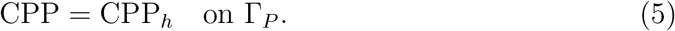
  – To model occlusions, blood flow rate through the superficial perfusion region associated with the occluded vessel is set to zero while perfusion pressure in other regions is maintained. For example, a right middle cerebral artery (R-MCA) occlusion is captured by

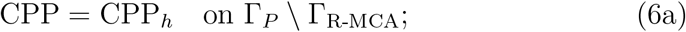

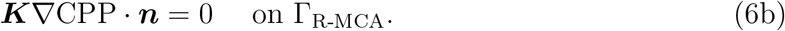

Once a boundary region Γ_x_ is defined, the corresponding volumetric blood flow rate Q_x_ [m^3^s^-1^] is computed as

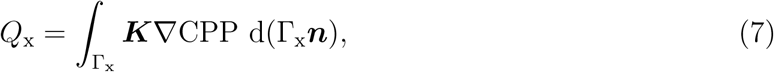

where the Darcy velocity vector is –***K***∇CPP so that *Q*_x_ is positive if blood flow is delivered to the brain. Setting Γ_x_ = *∂*Ω results in the total volumetric blood flow rate to the brain (*Q_t_*). Furthermore, Γ_x_ = Γ_L-MCA_ and Γ_x_ = Γ_R-MCA_ lead to *Q*_L-MCA_ and *Q*_R-MCA_, respectively. Thereafter, the representative volumetric blood flow rate of the middle cerebral arteries is obtained by averaging: *Q*_MCA_ = 0.5 · (*Q*_L-MCA_ + *Q*_R-MCA_). Volumetric blood flow rates corresponding to other major cerebral arteries are measured similarly. Alternatively, volumetric blood flow rates in major cerebral arteries can be estimated from voxelised CBF maps using a registered vascular territory atlas, e.g. [26]. In the case of simulations, errors originating from atlas registration and non-trivial CBF map voxelisation are avoided by utilising Equation (7) for volumetric flow rate measurements.

## 3 Methods

### 3.1 Participants

Data for 75 healthy control subjects (age 68 ± 8 years, range 47–86 years, 56% female) were drawn from the v500.0 release of the European Prevention of Alzheimer’s Dementia (EPAD) study (VUmc site only) [27]. Reference subjects with missing age or sex information, or imaging data were excluded from the baseline cohort. Both structural and perfusion-weighted images were obtained with a single 3T Achieva scanner (Philips Healthcare).

### 3.2 Image acquisition and processing

Structural images were measured using T1-weighted MRI with a spatial resolution of 1.0 × 1.0 × 1.2 mm^3^. CBF was measured with a pseudo-continuous ASL (PCASL) MRI sequence with a 2D echo-planar imaging (EPI) readout and the following parameters: repetition time/echo time (TR/TE) 4800/10.4 ms, labelling duration 1650 ms, voxel size 3.44 × 3.44 × 4.50 mm^3^ with 36 slices without a slice gap, and 30 control-label pairs. Post-labelling delay ranged between 2025 for the first slice and 3310 ms for the last slice. Two background suppression pulses were applied at 1710 ms and 3142 ms after the start of the labeling. A separate calibration M0 scan was acquired with the same acquisition protocol, but without background suppression or labeling, and with a TR of 10 s.

Neuroimaging data were processed using the ExploreASL MATLAB toolbox with default settings detailed in [28]. In short, T1-weighted images were segmented using CAT12 [29] to obtain probability maps of gray matter, white matter, and cerebrospinal fluid. The processing of ASL perfusion images included motion correction, motion outlier detection, and rigid-body registration of the CBF maps to the segmented gray matter images. CBF was quantified using a single-compartment model [30].

### 3.3 Computational perfusion model generation

The cerebral perfusion model is defined by a set of input parameters required to solve Equation (1) subject to the boundary condition Equations (4)-(6). The proposed computational perfusion model generation method integrates clinical data to set up simulations accordingly, as depicted in Figure 1.

**Figure 1:**
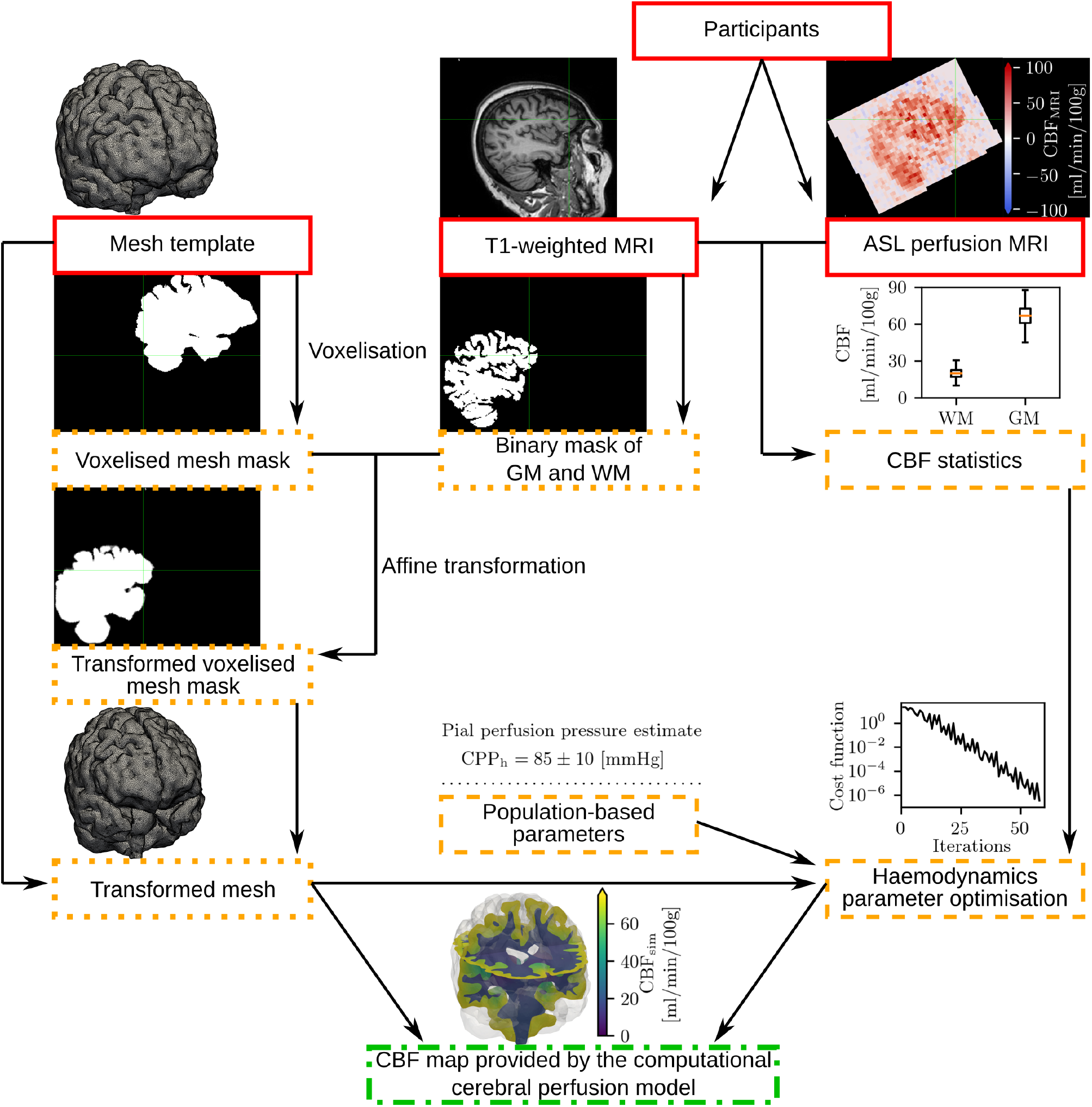
Schematic drawing of the computational perfusion model generation method. Solid boxes: input; dash-dotted box: output; dotted boxes: quasi-patient-specific brain mesh generation; dashed boxes: haemodynamic parameter tuning. ASL - arterial spin labelling, MRI - magnetic resonance imaging, CBF - cerebral blood flow, GM - grey matter, VM - white matter, CPP - cerebral perfusion pressure.

Currently, not every model parameter can be obtained in a patient-specific manner. Some of the necessary perfusion model parameters are available directly from clinical data (e.g. brain geometry). Other parameters are (i) inferred indirectly from clinical data (arteriole permeability); (ii) obtained based on population statistics (e.g. blood pressure), or (iii) inferred from pre-clinical measurements (e.g. grey matter coupling coefficient). For this reason, a computational model is not designed to mirror the properties of a specific individual at this stage. Instead, multiple models are introduced to be representative of the members of a patient cohort.

Computational perfusion model generation is subdivided into two sub-problems as shown in Figure 1:

1. Obtaining a tetrahedral mesh of brain geometry with labelled volume and surface regions as detailed in Section 3.4;
2. Inferring physiological parameters including the grey and white matter coupling coefficients (*c*^GM^ and *c*^WM^) and the arteriole permeability 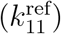 explained in Section 3.5.

### 3.4 Anatomically accurate brain geometry preparation

Perfusion simulations require a mesh including grey and white matter. Quasi-patient-specific geometries are obtained by first creating a general mesh template in the standard space and then an affine registration of this template to the tissue mask of each specific subject. This approach avoids time-consuming and frail patient-specific geometry reconstruction [31]. A mesh template is prepared from the averaged T1-weighted MNI template of the IXI555 cohort [29, 32], see Figures 2(a-c).

**Figure 2:**
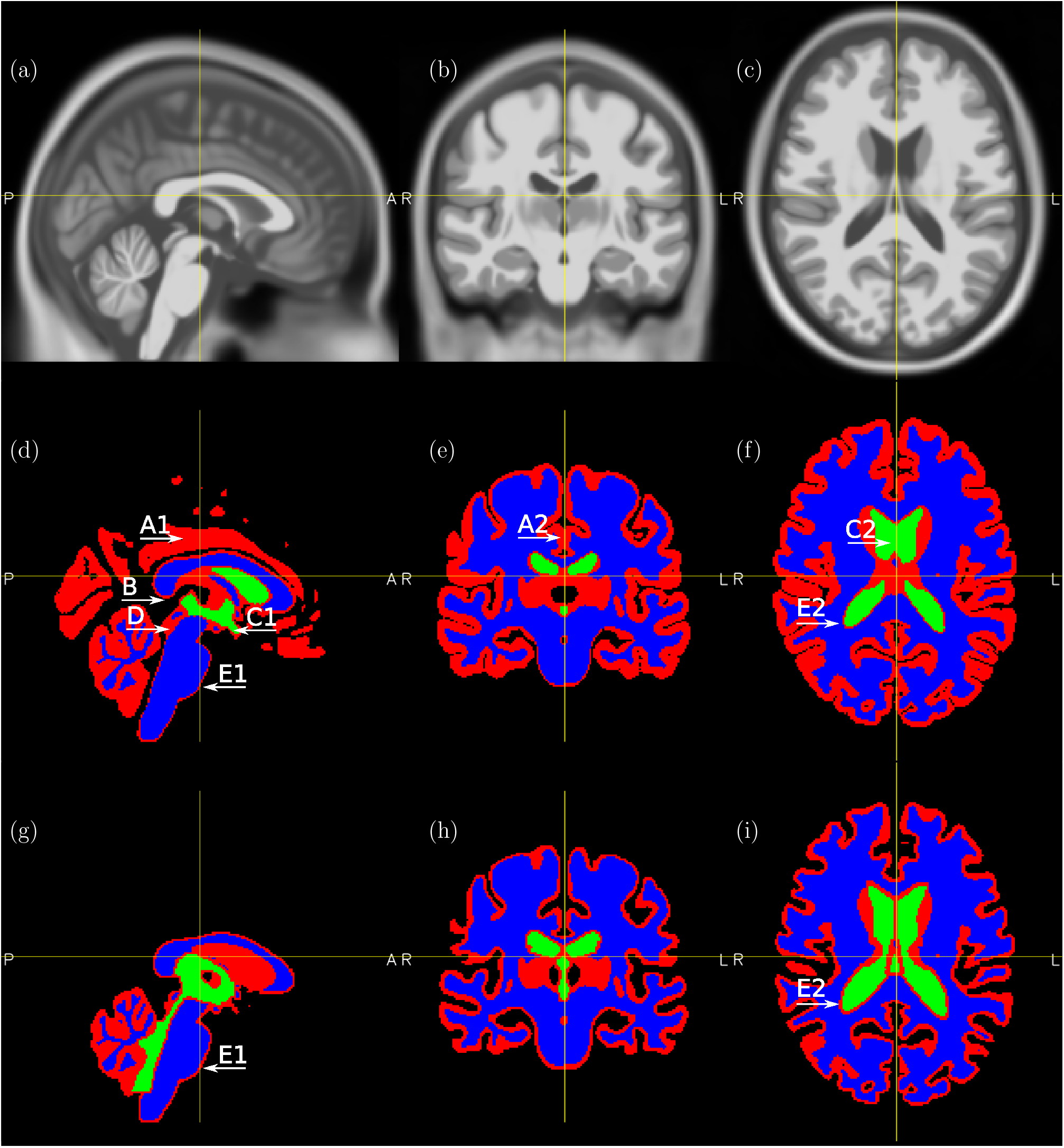
Visualisation of the T1-weighted MRI MNI atlas of the IXI555 cohort [29, 32] (a-c). Default SimNIBS [34] segmented images (d-f) and altered grey and white matter masks (g-i). Sagittal (a,d,g), coronal (b,e,h), and axial (c,f,i) slices. Colouring of (d-i): red - grey matter; blue - white matter; green - ventricles. The arrows indicate anatomical flaws in the segmentation: A - connected hemispheres; B - inaccurate opening between the subarachnoid space and the third ventricle; C - incomplete segmentation of ventricles; D - aqueduct of Sylvius labelled as grey matter; E - simplified tissue layering assumption.

First, the MNI template is segmented using the Anatomy Toolbox (CAT) [29, 33] to GM, WM, and CSF masks, see Figures 2(d-f). These masks have several issues originating from (i) the blurred averaged image, and (ii) the assumptions and simplifications in the segmentation and meshing algorithms:

A. The left and right hemispheres are not well-separated;
B. Connectivity between the ventricles and the subarachnoid space is inaccurate;
C. Connectivity between the lateral and third ventricles, and the surrounding tissue is inaccurate;
D. Connectivity between the third and fourth ventricles is inaccurate;
E. Connectivity of tissue and cerebrospinal fluid spaces is inaccurate (i.e. there is no direct connection between WM and CSF regions).

The automatically generated default mesh provided by SimNIBS (Mesh A) [34] is burdened by the same issues as shown in Figures 3(a-c).

**Figure 3:**
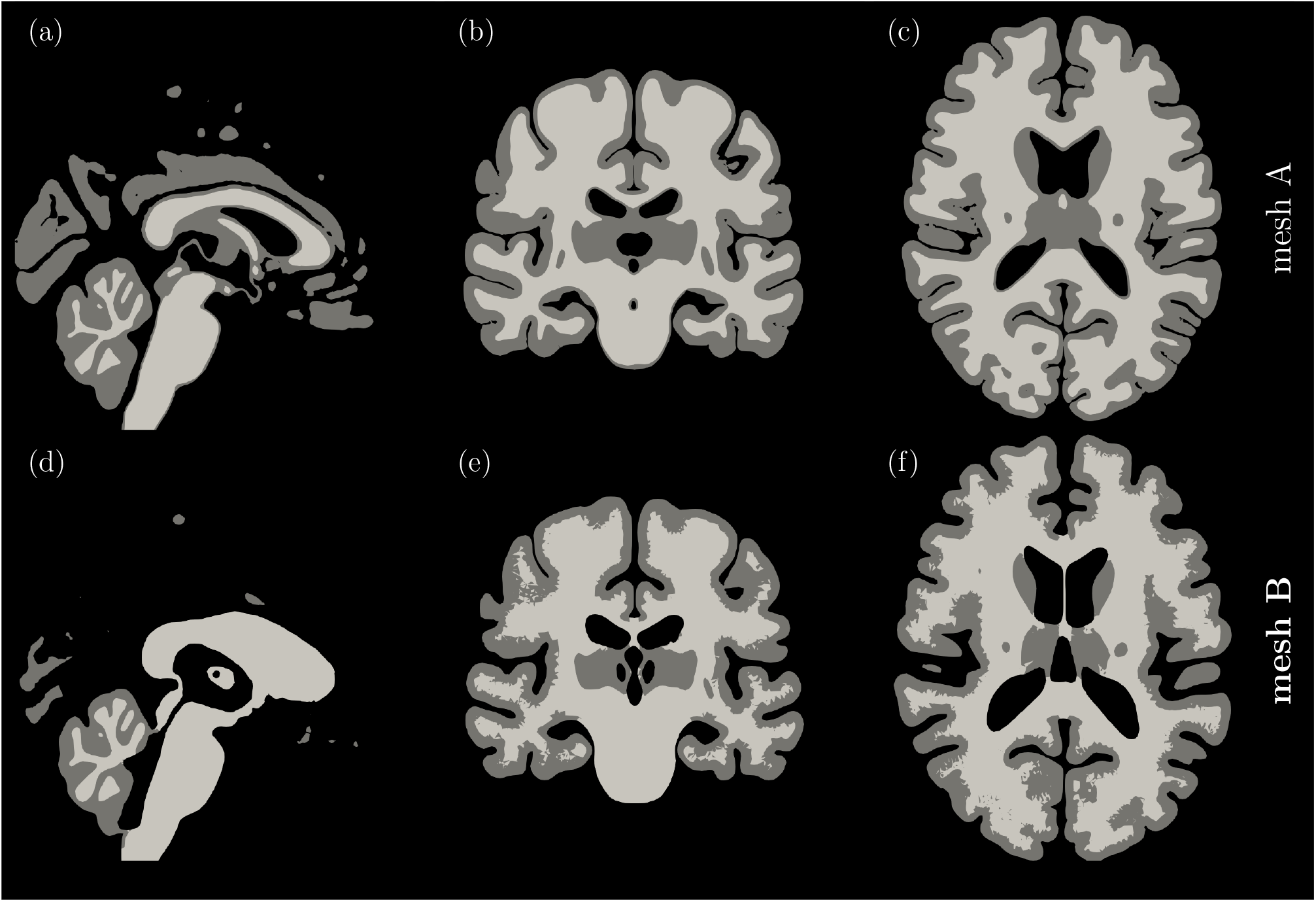
Visualisation of the grey and white matter domains of the SimNIBS [34] default Mesh A (a-c) and Mesh B (d-f) along sagittal (a,d), coronal (b,f), and axial (c,f) slices. Mesh B is chosen as the mesh template in Figure 1.

The majority of the listed issues (A-D) are addressed after segmentation by employing standard image processing algorithms and manual amendments. For example, the mask of the ventricles has been altered manually based on the original segmentation and the T1-weighted template, and the overall tissue mask has been eroded to ensure that anatomical domains are well-separated. (Ventricles are separated deliberately from the subarachnoid space to ease the imposition of boundary conditions.) Figures 2(g-i) depict the resulting adjusted segmentation.

The remaining issue E cannot be fixed at the segmentation stage, see Figure 2(g-i), as the SimNIBS mesh-generating algorithm relies strongly on a non-anatomical tissue layering assumption based on which a CSF layer is followed strictly by a GM and then a WM layer. The issue E is thus propagated to the mesh generated by SimNIBS and has to be corrected by post-processing the resulting mesh. Accordingly, tetrahedral GM elements are relabelled in the vicinity of the ventricles and the brain stem based on the original CAT [29, 33] segmentation registered to the mesh. Furthermore, tetrahedral WM elements are relabelled in the vicinity of the cortical surface to ensure that the corresponding GM thickness is at least ≈ 2 [mm] [35]. (A similar relabelling routine has been implemented previously in the Brain2Mesh toolbox [17] to address similar issues.) The resulting mesh is shown in Figures 3(a-c) and is referred to as Mesh B. Mesh B is selected as default for generating mesh template in the following text.

The next step is to generate a voxelised mesh template from Mesh B. Voxelisation is carried out using a ray intersection method [36]. Thereafter, a patient-specific T1-weighted MRI scan is segmented with CAT [29, 33] to obtain a binary tissue mask by thresholding GM and WM probability maps at 0.5.

The voxelised mesh mask is registered to the patient-specific tissue mask using an affine transformation with 12 degrees of freedom as implemented in the Linear Image Registration Tool (FLIRT) [37], part of the Oxford Centre for Functional MRI of the Brain (FMRIB) Software Library (FSL) [38]. The transformation is applied to the mesh template resulting in a quasi-patient-specific transformed mesh. The anatomical details of the computational models are analysed and compared to clinical measurements in Sections 4.1 and 5.1, respectively.

### 3.5 Physiological parameter optimisation

Psychological parameters of the virtual brain are obtained from real data and literature either as individual values or as population statistics.

Perfusion statistics are extracted from patient-specific T1-weighted and ASL MRI scans (Figure 1) based on the following quantities: (i) mean CBF in GM 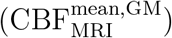; (ii) mean CBF in WM 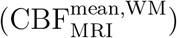; (iii) 20^th^ percentile of CBF 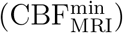; 80^th^ percentile of CBF 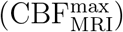. The percentile values are utilised as robust estimates of the minimum and maximum CBF, respectively. The “MRI” subscript distinguishes quantities deduced from structural/perfusion-weighted images from results corresponding to simulations (“sim” subscript).

Physiological parameter initialisation and optimisation based on the aforementioned perfusion statistics follows a similar strategy as in [13]. Population-based parameters are incorporated from the literature as detailed below.

1. Based on rodent experiments [39], pial perfusion pressure (CPP_*h*_) is well-approximated by the diastolic pressure. Therefore, the CPP_*h*_ pressure is selected randomly for each computational perfusion model based on diastolic pressure measurements in ischaemic stroke patients (Gaussian distribution assumed with mean and standard deviation as reported in [40]).
2. For a given 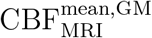 and CPP_*h*_ pair, the GM coupling coefficient is estimated as

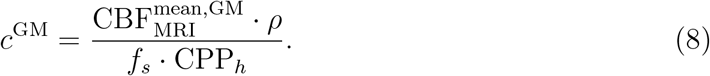 Here, *f_s_* is a scaling factor to represent the average arteriole perfusion pressure in the GM as a fraction of the boundary pressure (CPP_*h*_).
3. Parameters 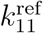 and *c*^WM^, are obtained by minimising the

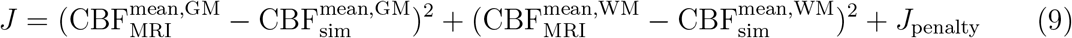

cost function. A penalty term (*J*_penalty_) ensures that CBF remains between the predefined minimum and maximum:

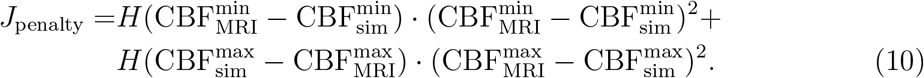 Here, *H* is the Heaviside function resulting in a non-zero *J*_penalty_ only if the 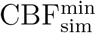 and 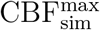 extrema are out of the target range determined by ASL MRI data.

Parameters are initialised based on a previous study [13] so that iterations start from a realistic perfusion map associated with a relatively low cost function value. Haemodynamics in the computational models in the healthy cases are analysed and compared to clinical measurements in Sections 4.2 and 5.2, respectively.

### 3.6 Acute ischaemic stroke simulations

Total occlusion of the middle cerebral artery is simulated in 75 virtual patients by setting blood flow rate through the boundary region corresponding to the single occluded vessel to zero, see Equation (6). This test case was chosen because middle cerebral artery occlusions are responsible for more than 60% of acute ischaemic stroke cases [41]. Both left and right hemisphere occlusions are simulated resulting in 150 virtual AIS patients. Two method are used to delineate the infarct volume based on the simulated CBF images:

1. IV_*r*_ – the region with a relative perfusion drop of 70%, see Figure 8(b) [42]. Disadvantage: overestimation of the ischaemic region compared to diffusion-weighted MRI [43].
2. IV_*a*_ – the region with an absolute perfusion value below 5 ml/min/100g. Disadvantage: a larger than 5 ml volume is labelled as infarct in 3 out of 75 virtual patients.

The simulated distribution of these proxies is evaluated and compared to clinical infarct measurements from the literature in Sections 4.3 and 5.2, respectively.

### 3.7 Numerical solution

Numerical solutions of the governing equations are obtained using the finite element method as implemented in the open-source FEniCS library [44]. The brain geometry is captured by tetrahedral elements. The weak form of Equation (1) is discretised using second-order Lagrange elements for *p*, and zeroth-order discontinuous Galerkin elements for the permeability tensor ***K*** and the coupling coefficient *c*.

For further details, we refer to [13, 20].

### 3.8 Statistical analysis

The anatomical and physiological fitness of the 75 computational perfusion models are investigated by means of linear regression analyses between variables measured in healthy reference subjects and brain models. The variables of interest selected for regression analyses are not used directly for the parameter tuning of the models. These variables include grey and white matter volumes (GMV and WMV), and the total volumetric blood flow rate to the brain (*Q_t_*). Pearson correlation coefficients (Pearson’s *r*) are computed between simualted and MRI samples utilising the “linregress” function of the SciPy [45] open-source Python library which relies on the P-value of the Wald test to evaluate statistical significance. The corresponding null hypothesis is that the slope of the linear regression model is zero which is rejected if the usual condition of *P* < 0.05 is met.

The anatomical fitness of Mesh A and mesh B are evaluated by calculating deviations from the EPAD dataset. The intracranial volume ratios of the brain tissue, GM, and WM are compared with mean values measured by the segmentation of healthy reference subjects’ T1-weighted images.

## 4 Results

### 4.1 Geometrical analysis of the transformed meshes

Figure 3 highlights several qualitative geometrical differences between Mesh A and Mesh B. A smooth grey-white matter interface cannot be preserved in Mesh B because of the relabelling required for preserving anatomical structures. Nevertheless, the grey white matter regions of Mesh B remain continuous with three separated regions corresponding to the two hemispheres and the cerebellum. Such qualitative anatomical details are not preserved by Mesh A. The tissue volume distributions in the computational models based on Mesh B and the T1-weighted MRI measurements of the EPAD cohort are in good agreement, Table 2. In the following, units are presented based on clinical conventions (e.g. volume in [ml], CBF in [ml/min/100g]) to ease direct comparison with existing datasets.

**Table 2:** Statistical analysis of the brain anatomy (geometry) and physiology (haemodynamics) of healthy control subjects and computational models (transformed Mesh B): SD - standard deviation; GMV - grey matter volume; WMV - white matter volume; Q_t_ - total volumetric blood flow rate to the brain.

| | Computational models<br>mean±SD | EPAD [27]<br>mean±SD | Pearson's $r$ | $P$-value |
| --- | --- | --- | --- | --- |
| GMV [ml] | $620 \pm 63$ | $593 \pm 49$ | 0.74 | $< 0.001$ |
| WMV [ml] | $542 \pm 55$ | $510 \pm 61$ | 0.84 | $< 0.001$ |
| $Q_t$ [ml/min] | $424 \pm 73$ | $463 \pm 84$ | 0.89 | $< 0.001$ |

Figure 4(a) highlights discrepancies between the tissue masks of models (transformed Mesh B) and healthy control subjects (EPAD), which are typically located near the boundary of the tissue geometry. The total brain volume of healthy controls tend to be overestimated by 58 ml (on average 5%) by the corresponding brain models, see Figure 4(b).

**Figure 4:**
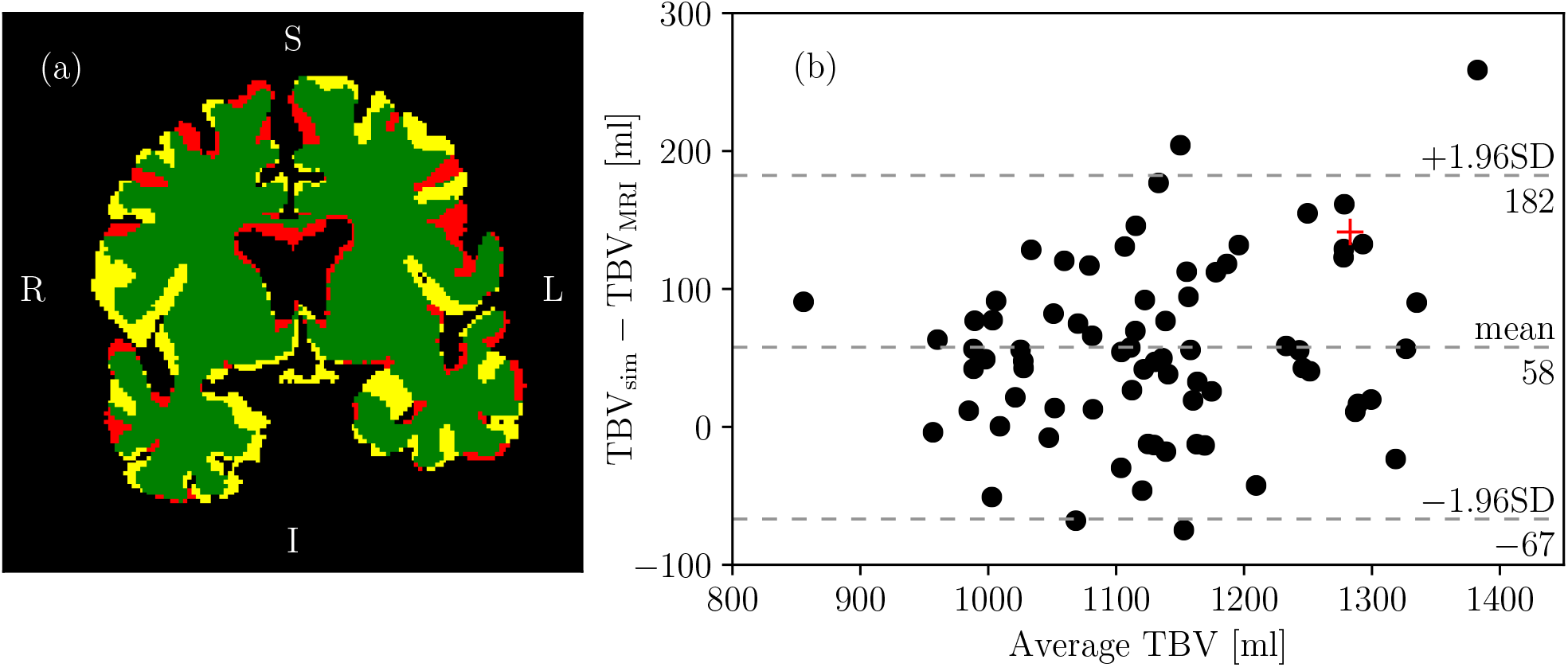
Comparison of the tissue mask corresponding to a healthy reference subject (sum of grey and white matter probability maps larger than 0.5) and the associated transformed voxelised mesh mask (a): green - overlap; yellow - reference only; red - perfusion model only. Bland-Altman plot comparing total brain volumes (TBV - sum of GM and WM) corresponding to reference subjects (TB_MRI_, measured as the sum of grey and white matter probability maps) and the associated quasi-patient-specific transformed mesh (TBV_mesh_) (b). The red + symbol in (b) marks the patient shown in (a). L - left, R - right, S - superior, I - inferior, SD - standard deviation.

### 4.2 Haemodynamic analysis of computational perfusion models

The cost function trends corresponding to the physiological parameter optimisation are displayed in Figure 5(a) and the resulting parameters are listed in Table 1. In about 57% of the cases, the square root of the cost function 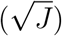 drops below one, meaning that the target grey and white matter CBF values are met within the specified minimum and maximum limits with a deviation less than 1 ml/min/100g. The optimisation leads to a final 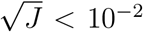 in 20 out of 75 cases. CBF values displayed in Figures 5(b-c) indicate that the simulations overshoot regional target values in every case. In the worst-case distinguished in Figures 5(a-c), 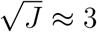 ml/min/100g is reached. This relatively large difference is caused by a mismatch between the targeted and the reached average grey matter perfusion values as suggested by Figure 5(b).

**Figure 5:**
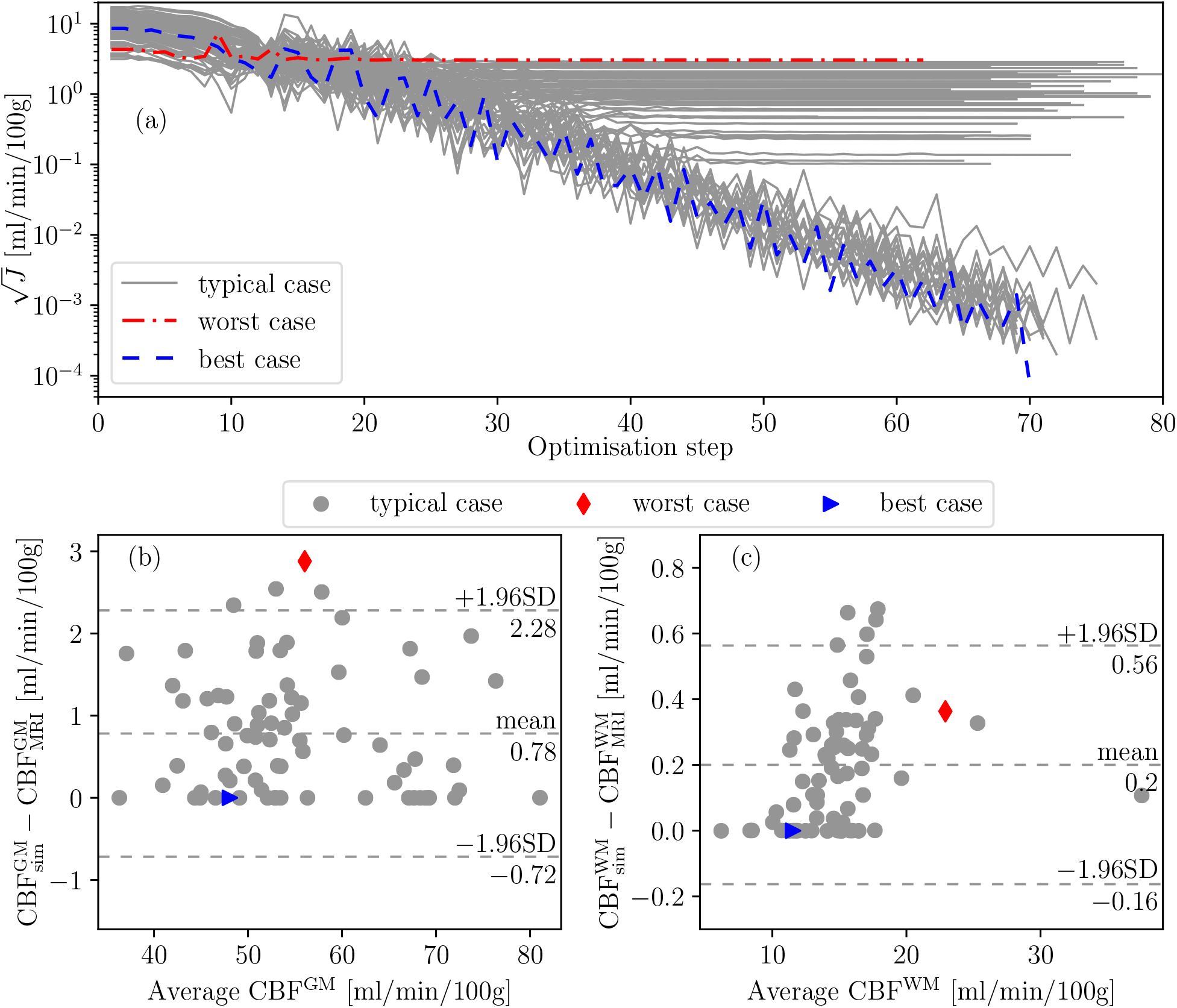
Cost function (*J*) convergence trends as functions of the optimisation steps (a); Bland-Altman plots of the mean white matter perfusion CBF^mean,WM^ (b) and mean grey matter perfusion CBF^mean,GM^ (c) of computational brains and healthy references. Worst and best cases correspond to the highest and lowest final cost function values, respectively. The subscripts “sim” and “MRI” distinguish data corresponding to simulations and ASL perfusion MRI, respectively. CBF - cerebral blood flow, SD - standard deviation.

Considering total volumetric blood flow rate to the brain (*Q_t_*), virtual cases show a good agreement with values inferred from ASL perfusion MRI of healthy reference individuals in EPAD as detailed in Table 2. Volumetric blood flow rates in computational brain models tend to be underestimated on average by 8%. The average CBF in perfusion models is 37 ml/min/100g whereas the average CBF of healthy references in the EPAD cohort is 42 ml/min/100g based on the corresponding ASL MRI measurements. Whereas average grey and white matter perfusions are matched by the optimisation algorithm, computational models underestimate total brain perfusion on average by 12%. This relatively high error originates from the overestimation of the brain volume and the underestimation of the total volumetric blood flow rate. Nevertheless, Pearson’s *r* and the *P*-value presented in Table 2 emphasise a strong, statistically significant correlation between the total blood flow rates measured in healthy reference subjects and in the computational cerebral perfusion models.

Figures 6(a-c) and (d-f) visualise 3D CBF maps corresponding to a clinical measurement and the linked brain model. Perfusion distribution in the computational model appears to be relatively homogeneous compared to clinical measurements. The perfusion model parameters could thus be tuned further to achieve better agreement with clinical data representing instantaneous or short-term averaged perfusion state. Instead, we recommend interpreting simulation results as time-averaged measurements which are reminiscent of clinical studies reporting ensemble-averaged perfusion maps [46].

**Figure 6:**
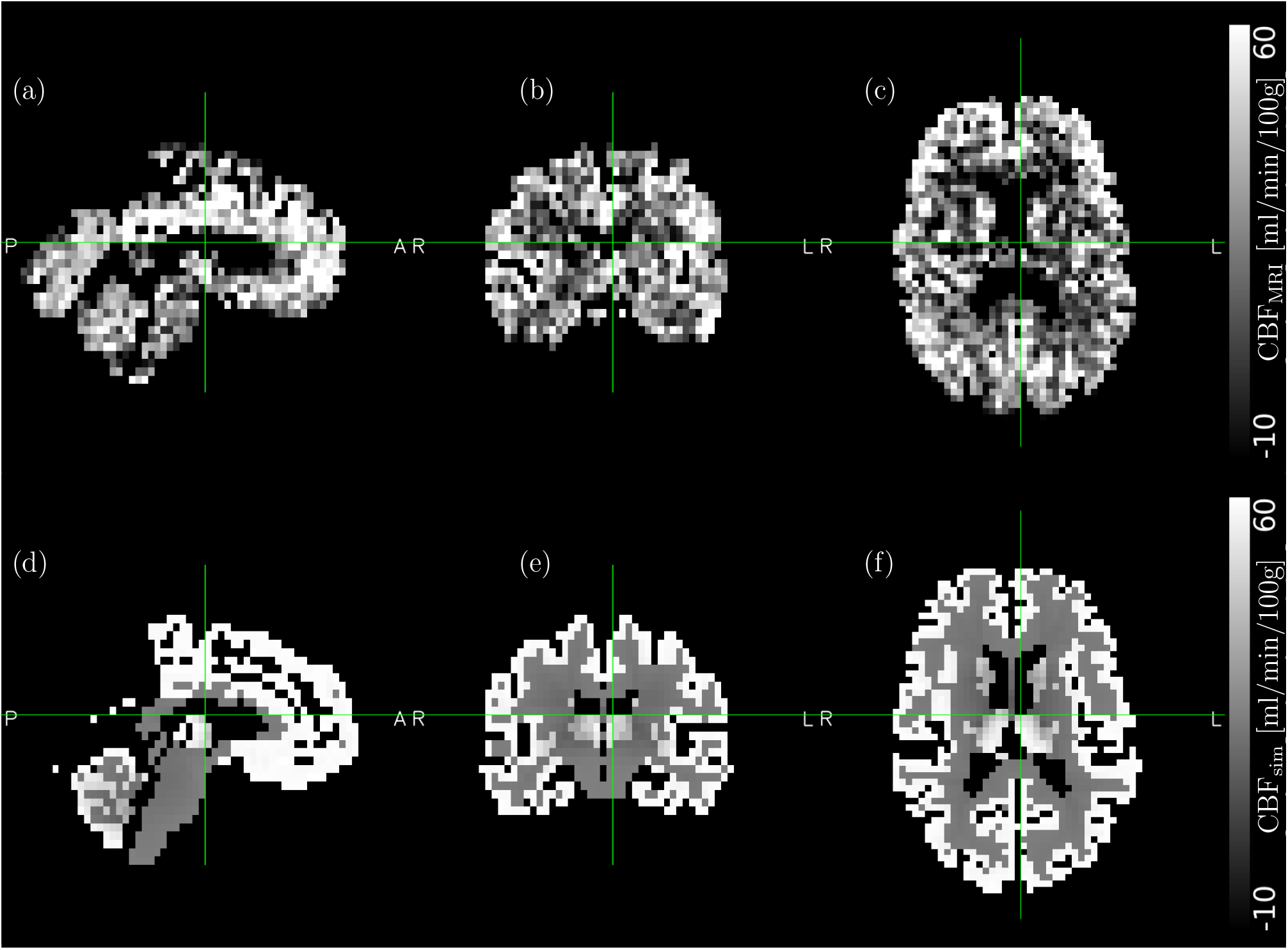
Cerebral blood flow (CBF) maps of a healthy reference individual from the EPAD project (a-c) and the corresponding simulation (d-f) along sagittal (a,d), coronal (b,f), and axial (c,f) slices. In the case if (a-c), MRI results are multiplied by a binary tissue mask which corresponds to regions where the sum of the grey and white matter probability maps is higher than 0.5. The subscripts “sim” and “MRI” distinguish data corresponding to simulations and ASL perfusion MRI, respectively. Simulation data linked to the worst-case highlighted in Figure 5.

The probability density functions displayed in Figure 7 confirm that the simulated grey and white matter regions have well-distinguishable perfusion values. This is also visible in Figures 6(d-f). By comparison, the probability density function of ASL perfusion MRI shows a nearly Gaussian trend with standard deviation in the same order of magnitude as its mean value. This is probably caused by (i) the relatively low signal to noise ratio of ASL perfusion MRI, and (ii) the continuous fluctuations of local cerebral perfusion associated, e.g. with neurovascular coupling.

**Figure 7:**
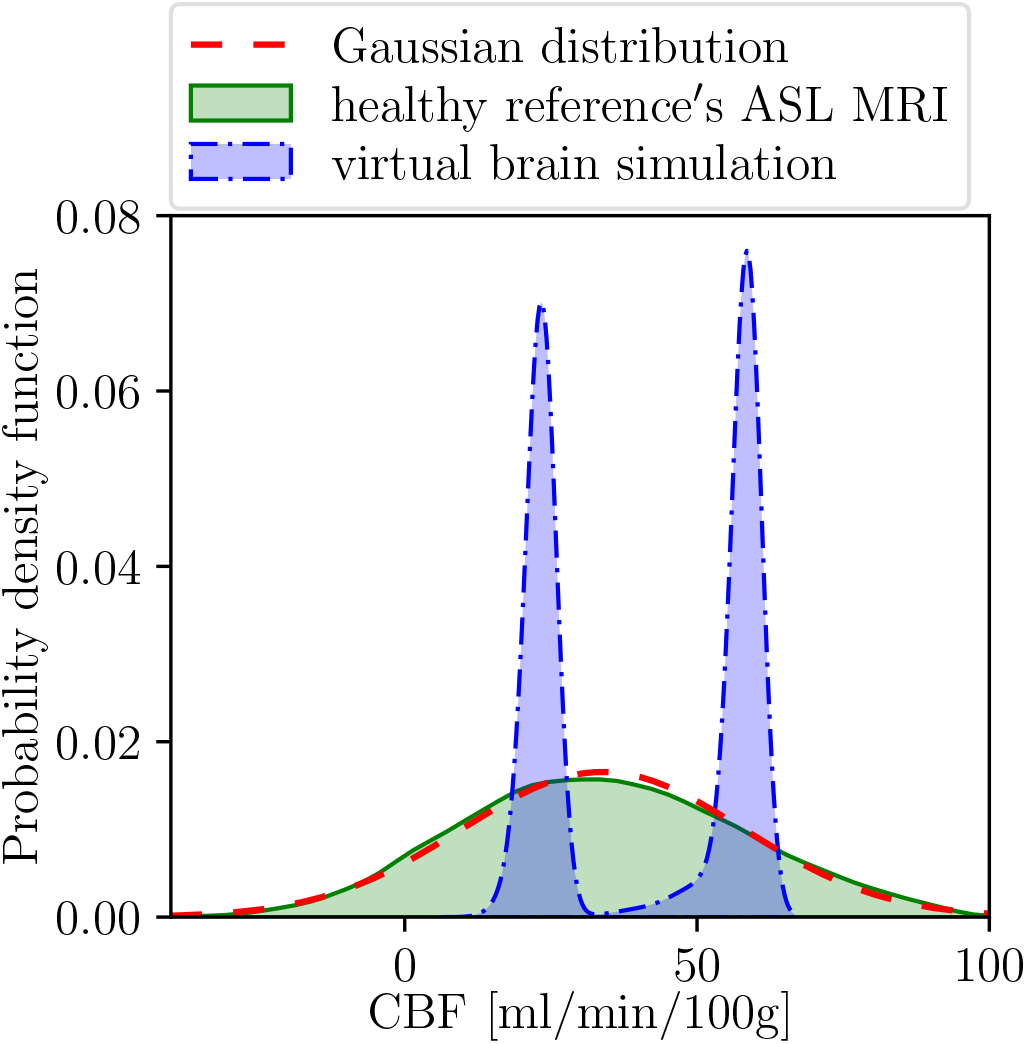
Probability density functions of cerebral blood flow (CBF) maps. The subscripts “sim” and “MRI” distinguish data corresponding to simulations and arterial spin labelling (ASL) perfusion MRI, respectively. MRI results are multiplied by a binary tissue mask labelling regions where the sum of the grey and white matter probability maps is higher than 0.5. The perfusion maps of a healthy reference subject providing the input for this graph are displayed in Figure 6. The distribution of CBF_MRI_ data is well approximated by the Gaussian function (mean±SD: 34±24 ml/min/100g). SD - standard deviation.

### 4.3 Infarct volume estimation in acute ischaemic stroke

Perfusion pressure and perfusion distributions in a typical occluded scenario are visualised in Figures 8(a) and (b), respectively. Both infarct proxies result in similar, approximately normally distributed, infarct volume estimates. Mean and standard deviation values in the case of the relative perfusion drop based infarct volume proxy (IV_*r*_) are 205 ± 23 ml. By comparison, IV_*a*_ leads to somewhat smaller infarct estimations with higher variance: 197 ± 25 ml.

**Figure 8:**
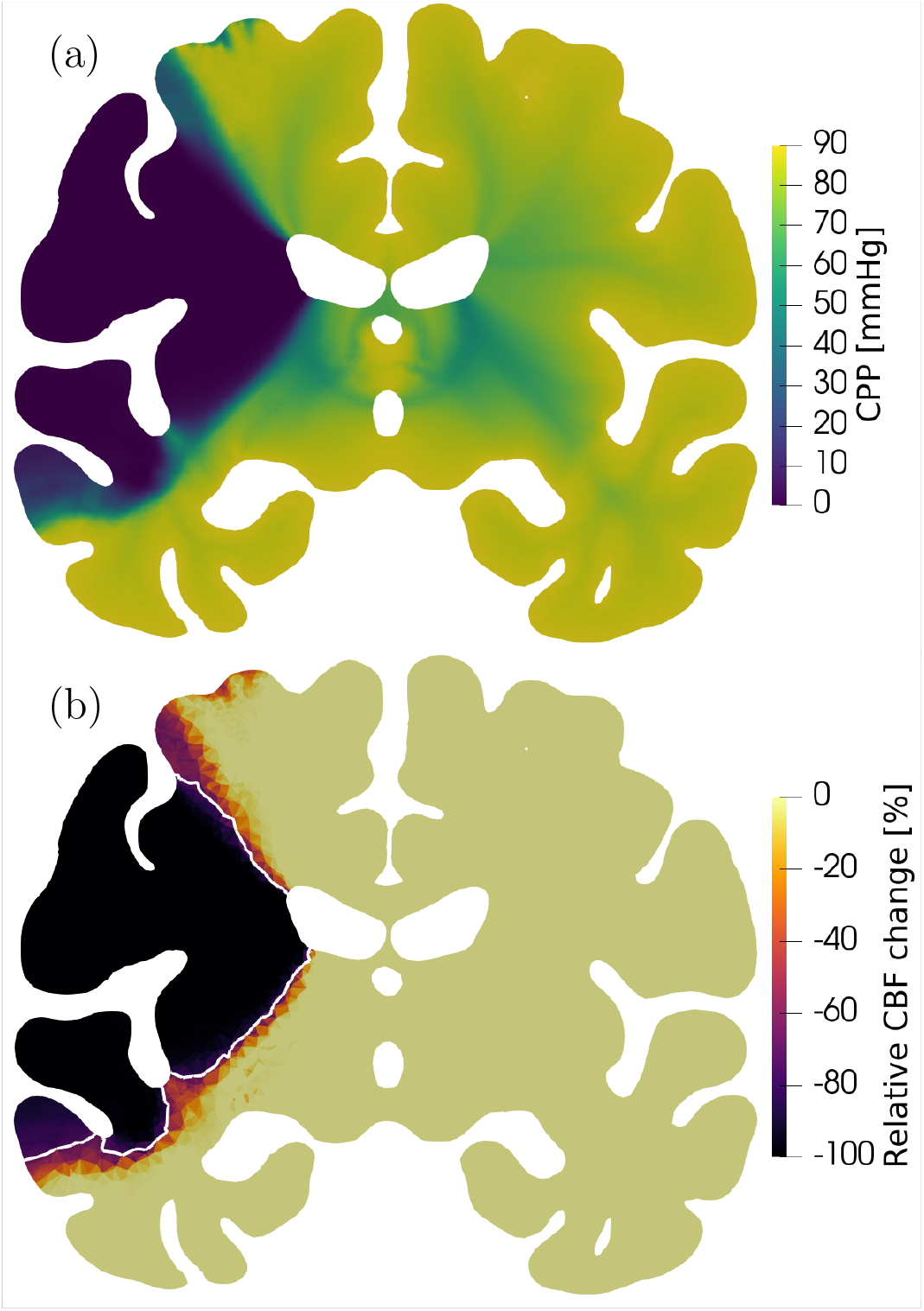
Simulation results in an occluded case corresponding to the virtual patient shown in 6(d-f): CPP - cerebral perfusion pressure (a) and relative cerebral blood flow (CBF) change in addition to isolines indicating 70% CBF drop used to define IV_*r*_, the infarct volume based on relative CBF change (b).

## 5 Discussion

The proposed computational cerebral perfusion model generation approach integrates human neuroimaging data to infer anatomical (geometrical) and haemodynamic model parameters, and thus reach the goal of the present study regarding the realisation of cohort-level perfusion simulations. The present study reports overall 225 brain simulations including 75 healthy cases, and 150 ischaemic stroke cases with 75 left and 75 right middle cerebral artery occlusions. Group-level tissue volume and cerebral blood flow measurements in the model brains match clinical observations corresponding to the healthy reference cohort used for model parameter tuning. Furthermore, the model is capable of providing predictions in cohorts of acute ischaemic stroke patients.

In this section, the proposed perfusion-focused computational brain generation approach is evaluated. Simulations and the utilised clinical data are analysed in comparison with independent datasets. The anatomical and physiological properties of interest are listed in Table 3 alongside reference values from the literature.

**Table 3:** Comparison of simulations and clinical measurements corresponding to healthy reference subjects and to reference data from the literature. Mean ± standard deviation values are shown as applicable. Anatomical variables include TBV - total brain volume (sum of GM and WM); GMV - grey matter volume; WMV - white matter volume; ICV - intracranial volume; *A*_total_ - total brain surface area; *A*_cortical_ - cortical surface area. Physiological variables include volumetric blood flow rates corresponding to the entire brain (*Q_t_*), and the anterior *Q*_ACA_, middle *Q*_MCA_, and posterior *Q*_PCA_ cerebral arteries. Reference data are gathered from the following sources: (i) Nielsen *et al.* [34]*: default SimNIBS head model (mesh A); (ii) Huang *et al.* [16]^†^: NYHM - New York Head Model, age range 18.5-43.5; (iii) Ikram *et al.* [47]^‡^: T1-weighted and T2-weighted MRI, 490 subjects, 249 women, 73 ± 8 years old; (iv) Lüders *et al.* [23]^§^: T1-weighted MRI, 100 subjects, 50 women, 25 ± 4 years old; (v) Winkler *et al.* [63]^¶^: T1-weighted MRI-based surface reconstruction; (vi) Zarrinkoob *et al.* [55]^||^: phase-contrast (PC) MRI, 45 subjects, 23 women, 71 ± 4 years old. (vii) Xing *et al.* [54]^**^: PC MRI, 39 subjects, 24 women, 71 ± 4 years old.

|  | Brain models | EPAD [27] | Reference data |  |
| --- | --- | --- | --- | --- |
|  |  |  | value | source |
| TBV/ICV [%] | 71.4 | $74.1 \pm 4.0$ | 78.8 | mesh A [34]* |
|  |  |  | 83.3 | NYHM [16]<sup>†</sup> |
| | | | $77.4 \pm 3.7$ | MRI [47]<sup>‡</sup> |
| GMV/ICV [%] | 38.1 | $39.9 \pm 2.6$ | 41.9 | mesh A [34]* |
|  |  |  | 49.2 | NYHM [16]<sup>†</sup> |
| | | | $55.1 \pm 2.0$ | MRI [23]<sup>§</sup> |
| | | | $46.6 \pm 4.0$ | MRI [47]<sup>‡</sup> |
| WMV/ICV [%] | 33.3 | $34.2 \pm 2.2$ | 36.9 | mesh A [34]* |
|  |  |  | 34.1 | NYHM [16]<sup>†</sup> |
| | | | $27.4 \pm 2.0$ | MRI [23]<sup>§</sup> |
| | | | $30.8 \pm 5.7$ | MRI [47]<sup>‡</sup> |
| $A_{\text{total}}$ [cm<sup>2</sup>] | 2040 | — | 2262 | mesh A [34]* |
|  |  |  | 1774 | NYHM [16]<sup>†</sup> |
| $A_{\text{cortical}}$ [cm<sup>2</sup>] | 1659 | — | 1770 | MRI [63]<sup>¶</sup> |
| $Q_t$ [ml/min] | $424 \pm 73$ | $463 \pm 84$ | $657 \pm 94$ | PC MRI [55] |
| | | | $488 \pm 123$ | PC MRI [54]** |
| $Q_{\text{ACA}}$ [ml/min] | $48 \pm 8$ | — | $75 \pm 15$ | PC MRI [55] |
| $Q_{\text{ACA}}/Q_t$ [%] | $11 \pm 2$ | — | $11 \pm 2$ | |
| $Q_{\text{MCA}}$ [ml/min] | $85 \pm 15$ | — | $131 \pm 23$ | |
| $Q_{\text{MCA}}/Q_t$ [%] | $20 \pm 4$ | — | $20 \pm 4$ | |
| $Q_{\text{PCA}}$ [ml/min] | $47 \pm 8$ | — | $51 \pm 10$ | |
| $Q_{\text{PCA}}/Q_t$ [%] | $11 \pm 2$ | — | $8 \pm 2$ | |

### 5.1 Anatomical evaluation

Obtaining computational geometries for physiological simulations is a non-trivial task. In the present study, we propose the creation of a mesh template as an intermediate step towards the standardisation of computational geometries. Thereafter, quasi-patient-specific geometries can be obtained by rapid and robust transformations (i.e. affine) instead of relying on resource-intensive, and fragile mesh generation algorithms. The present approach is suitable to produce a population of computational cerebral perfusion models with brain volume variations representative of patient cohorts used in clinical trials as detailed in Tables 2 and 3.

The mesh template plays a critical role in the present cohort-level modelling approach because any issues of the mesh template are naturally “inherited” by the generated brain models. The first half of Table 3 aims to evaluate the utilised mesh template (Mesh B) compared to other head models including Mesh A [34] and the NYHM [16]. The ratios of GM and WM volumes to ICV of Mesh A and Mesh B are representative of the EPAD healthy elderly population. The root mean square error of the GMV/ICV and WMV/ICV ratios of Mesh A and the NYHM compared to the EPAD mean value measurements is 2.4% and 6.6%, respectively. By comparison, the total error of Mesh B is 1.4% mirroring the superiority of Mesh B as a mesh template for elderly populations.

The TBV/ICV, GMV/ICV and WMV/ICV tissue volume ratios of the computational brains and the EPAD cohorts are lower than the reported values in previous neuroimaging studies [23, 47]. The differences are obvious especially when data from the present study are compared with values reported in the MRI study of Lüders *et al.* [23] and with measurements of the NYHM [16]. It is important to acknowledge that these datasets correspond to healthy young reference subjects (details in the caption of Table 3). Therefore larger TBV/ICV, GMV/ICV, and WMV/ICV tissue volume ratios are not surprising compared to the present investigation which focuses on healthy elderly references. The results highlight that it might be necessary to utilise different mesh templates for the perfusion simulations of cohorts with different age groups.

To date, only a few studies considered physiological simulations of human brains at the level of patient cohorts [48]. State-of-the-art segmentation [29, 49–51] and mesh generation techniques [17, 34, 48] are often computationally more expensive than all the other steps shown in Figure 1 and they have not been designed to satisfy requirements for perfusion simulations. The proposed perfusion model generation method gives up on patient-specific mesh generation to enable cohort-level organ-scale cerebral perfusion simulations.

Publicly available head or brain meshes reflect well the challenges associated with automated, anatomically accurate patient-specific mesh generation. For example, the grey and white matter regions of Mesh A [34], the NYHM [16] and the finite element mesh of Garcia-Gonzalez *et al.* [52] are simply unsuitable for perfusion simulations because of qualitative anatomical flaws. These meshes are burdened by segmentation artefacts leading to non-anatomical connections between the left and right hemispheres as shown in Figures 2(d-f) and 3(a-c). The aforementioned head models were designed for eletrophysiological and tissue deformation simulations where such details are less important. However in haemodynamics simulations, these artefacts manifest into non-physiological blood flow pathways.

### 5.2 Physiological evaluation in health and disease

The perfusion simulator integrates information extracted from patient-specific perfusion images to create computational brain models with brain perfusion statistics similar to measurements in healthy reference subjects. Although the individual signal to noise ratio of ASL MRI in WM is relatively low [30], it does not undermine testing the proposed perfusion model generation concept. The resulting computational model utilises the simplest formulation that can produce results reminiscent of perfusion maps measured in clinical scenarios [46, 53].

The second half of Table 3 presents a comparison of blood flow rate information obtained from simulations and clinical neuroimaging datasets. The *Q_t_* distribution resulting from simulations and the EPAD ASL perfusion MRI measurement agree well with previous clinical PC MRI measurements [54]. Conversely, the simulated blood flow rates in major cerebral arteries are significantly lower than reported previously [55]. However, normalised blood flow rates in major cerebral arteries (e.g. *Q*_MCA_/*Q_t_*) are in satisfactory agreement with previous measurements [55].

The context of use of the presented brain model generation method is to evaluate the impact of various therapies on cerebral perfusion which drives infarct formation. The present simulations do not account for compensatory mechanisms – such as blood flow rerouting through collaterals [2, 56] and autoregulation [57] – which could help to restore perfusion levels after major cerebral artery blockage. In addition, simulation results are expected to represent worst-case scenarios with relatively large infarct volume estimates because of the complete blockage of the middle cerebral artery. Therefore, clinical measurements corresponding to patients with MCA occlusion whose outcomes were “dependent” or “dead” after 3 months are selected from the literature [58] for a comparison shown in Figure 9. Real patients’ lesions have higher variance and the simulated worst-case scenarios can explain only half of the clinically measured infarct volume distribution [58]. In particular, simulations capture solely the largest infarcts ranging from about 140 to 275 ml whereas the range of clinical measurements is approximately [5; 275] ml.

**Figure 9:**
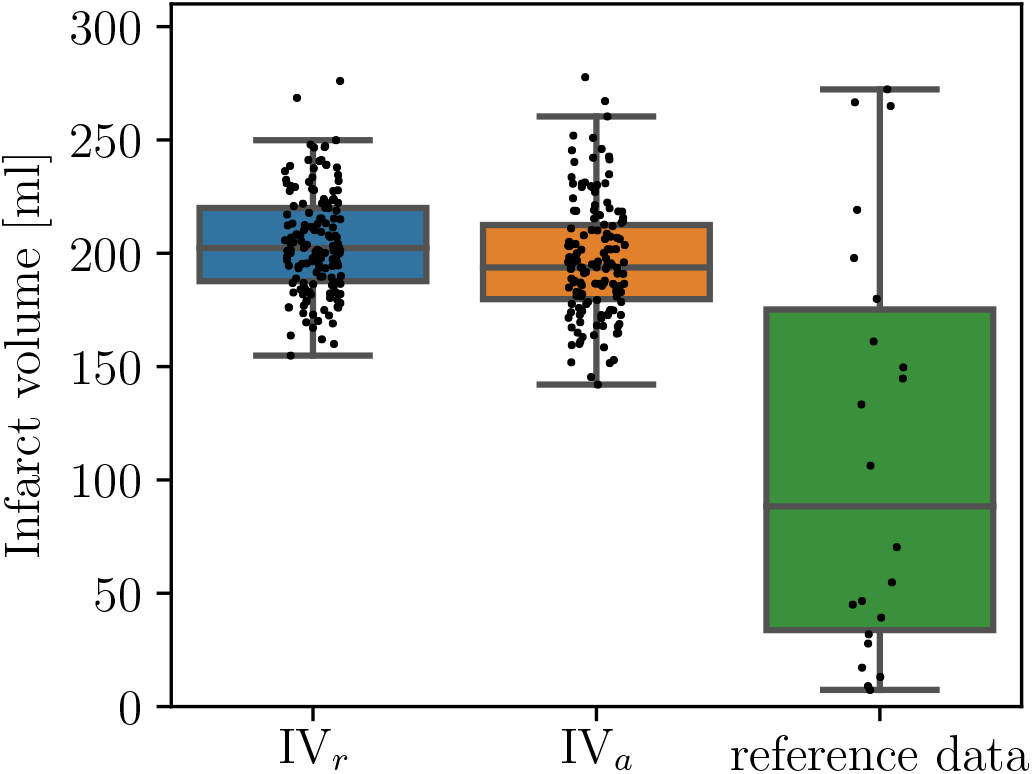
Comparison of infarct volume proxies defined based on relative cerebral blood flow (CBF) change (IV_*r*_) and from the absolute CBF level (IV_*a*_). Required CBF maps determined by middle cerebral artery occlusion simulations. The reference data [58] correspond to T2-weighted magnetic resonance imaging of acute ischaemic stroke patients with middle cerebral artery occlusions.

The present study demonstrates how cerebral perfusion simulations can integrate existing knowledge from various sources, including pre-clinical and clinical investigations. The results further computational perfusion modelling as a novel tool to boost medical device and drug development in a virtual environment by breaking previous barriers associated with small sample size.

### 5.3 Limitations

This study has several limitations. First, the affine transformation of the mesh template preserves the ratios of various tissue regions. Therefore, variations in grey and white matter ratios cannot be captured by the proposed computational cerebral perfusion model generation approach. It requires further investigations to evaluate whether nonlinear mesh transformation is suitable to capture more subtle anatomical details (e.g. variations in the volume fraction of tissue regions and the shape of ventricles).

Second, beyond anatomical factors, simulations depend on a set of physiological parameters characterising the spatial distribution of vascular resistance inside the brain tissue. For example, the vector space aligned with the centrelines of descending arterioles is necessary to compute the permeability tensor ***K*** representing the conductance of the corresponding blood vessels. However, experimental techniques have not yet been proposed to determine these crucial parameters.

Third, the test case regarding occluded scenarios emphasises the importance of crucial compensatory mechanisms, e.g. collateralisation [2, 56], which lowers the chance of infarct formation. Future development should aim to incorporate such effects into computational models as proposed in [59, 60]. Without such details, simulations cover solely the ‘right tail’ of the infarct volume distribution representing the largest infarcts. Furthermore, it is important to add processes which can be responsible for the lack of reperfusion after recanalisation, such as pericyte constriction [61] and microembolism [7, 62] to account for treatment effects. The present modelling framework does not incorporate such effects, but it does provide a solid foundation to facilitate such future extensions in cohort-level perfusion simulations.

## 6 Conclusions

The present study investigated an automated computational brain model generation approach which overcomes sample size limitations associated with perfusion simulations. The brain model generation method accounts for anatomical (geometrical) and physiological (haemodynamic) variations by integrating T1-weighted and arterial spin labelling MRI for parameter tuning. Whereas model parameters are tuned based on 75 healthy reference subjects, model predictions corresponding to cases with middle cerebral artery occlusions are validated against data in acute ischaemic stroke patients. The proposed method thus allows us to conduct perfusion simulations with significant anatomical and physiological variations in healthy and in ischaemic stroke cases. Simulations provided qualitatively and quantitatively realistic predictions regarding the infarct location and size. The results highlight that perfusion models in their current state can capture solely the largest infarcts because of the lack of compensatory pathways (e.g. collaterals) which could be added in future studies.

## Acknowledgements

This work was supported by the European Union’s Horizon 2020 research and innovation program, the INSIST project, under grant agreement No 777072. TIJ and SJP are grateful to the members of the INSIST consortium for useful discussions and their sustained support. TIJ would like to thank Giulia Luraghi of the Politecnico di Milano and Kevin Mattheus Moerman of National University of Ireland Galway for the valuable discussions on brain mesh generation. The EPAD data were obtained with support from the following EU/EFPIA Innovative Medicines Initiatives (1 and 2) Joint Undertakings: EPAD grant no. 115736, AMYPAD grant no. 115952. HJMMM was supported by the Dutch Heart Foundation (2020T049), and by the Eurostars-2 joint program with co-funding from the European Union Horizon 2020 research and innovation program, provided by the Netherlands Enterprise Agency (RvO).

## Authors’ contributions

TIJ carried out the research and prepared a draft of the manuscript under the supervision of SJP (mathematical and computational model development) and HJMMM (clinical data processing). JP and FB provided additional guidance to ensure the efficient usage of the ExploreASL image processing pipeline and the EPAD data set, respectively. Every author contributed to the revision of the paper.

